# Comparative phosphoproteome analysis of primary and metastatic uveal melanoma cell lines

**DOI:** 10.1101/2023.12.06.570407

**Authors:** K. Glinkina, R. Gonzalez Prieto, A.F.A.S. Teunisse, M.C. Gelmi, M.J. Jager, A.C.O. Vertegaal, A.G. Jochemsen

**Affiliations:** Department of Cell and Chemical Biology, Leiden University Medical Center, Leiden, The Netherlands; Department of Ophthalmology, Leiden University Medical Center, Leiden, The Netherlands

**Author notes:** Corresponding author: Oncogenic Signaling in Melanoma group, Department of Cell and Chemical Biology, Leiden University Medical Center, Postbus 9600, 2300 RC Leiden, The Netherlands.

**Keywords:** Uveal melanoma, metastases, proteomics, mass-spectrometry, MARK3, Rho GTPase

## Abstract

Uveal melanoma (UM) is an ocular tumor that often develops asymptomatically. Statistically, every second patient eventually develops metastases that drastically worsen the prognosis by several months of overall survival. While isolated liver perfusion with melphalan and more recently immunotherapy (Tebentafusp) are the few treatment options available for metastatic UM patients, their application is complex or expensive. There is an urgent need to understand drug response and identify potential avenues for therapy. Hence, we focused on uncovering altered phosphorylation signaling events in metastatic UM using proteomics as an approach to identify potential drug targets.

We analyzed the phosphoproteomes of the primary UM cell line Mel270 and two cell lines OMM2.3 and OMM2.5, derived from metastatic lesions of the same patient. We found 177 phosphosites to be altered significantly between primary and metastatic cell lines. Pathway analysis of up-regulated phosphosites in metastatic lines suggests that Rho signaling and mitotic cell cycle to be significantly altered uncovering potential routes of signaling for metastasis. Clinical data from LUMC and TCGA datasets uncovered *MARK3* expression (which links to Rho signaling) correlation with chromosome 3 status, a prognostic marker in UM, suggesting that MARK3 kinase might be involved in metastatic UM signaling.

## Introduction

Uveal melanoma (UM) is a rare tumor originating from the melanocytes located in uveal tract of the eye. In many cases UM develops asymptomatically and metastasizes by the time of the diagnosis [1]. Spread of UM metastases affects up to 50% of the patients and drastically worsen the prognosis due to resistance of the metastatic lesions to commonly used therapeutics [2]. Several therapeutic options available for metastatic UM patients include isolated liver perfusion with melphalan and recently developed immunotherapy approach (Tebentafusp) [3–4]. However, these treatment options are suitable only for subsets of patients, are complex or expensive. Therefore, there is an urgent need to understand drug response and identify novel avenues for therapy.

The genetic profile of UM is distinct from one of cutaneous melanoma. UM lacks the mutations in *BRAF* and *NRAS*, common for cutaneous melanoma; instead, virtually all UM cases are characterized by activation of Gα-proteins signaling cascade. In more than 90% of UM cases the mutations are harbored in *GNAQ* and *GNA11* genes, encoding Gαq and Gα11 subunits respectively; the remaining cases are characterized by the activating mutations in G-protein coupled receptor *CYSLTR2* or in signal mediator *PLCB4* [5–7]. The persistent activity of the Gα-proteins signaling cascade dysregulates several downstream pathways [8–10]. It fuels in activity of RhoA and downstream effectors, which trigger translocation to nucleous of YAP1 and TAZ, start YAP1-dependent transcription and initiation of malignant transformation of uveal melanocytes [11–14].

Secondary somatic alterations most commonly occur in the genes *EIF1AX, SF3B1*, *BAP1* in a mutually exclusive manner [15–17]. The presence of inactivating mutations in the translation initiation factor EIF1AX correlates with disomy of chromosome 3 and more favorable prognosis, while inactivation of splicing modulator SF3B1 is associated with intermediate metastatic risk and late-onset metastases. Inactivating mutation in *BAP1* gene followed by loss of chromosome 3 leads to complete depletion of BAP1 expression and strongly correlates with metastases development and poor prognosis. Besides the somatic mutations, copy number alterations on the chromosomes 1q, 3, 6p, 6q, 8q, 16q are common in metastatic UM [18–20].

Despite recent progress in unraveling the genetic mechanisms of UM, limited studies are available examining the (phospho)proteome of primary UM and metastatic UM [21, 22]. In this study we analyzed the phosphoproteome of the primary UM cell line Mel270 and the cell lines OMM2.3 and OMM2.5, derived from metastatic lesions of the same patient. We found 177 differentially phosphorylated sites between primary tumor and metastases, identified up- and down-regulated signaling cascades in metastases and suggested that MARK3 kinase might be involved in YAP1/TAZ signaling regulation.

## Materials and Methods

### Cell culture

Cell lines Mel270 (CVCL_C302), OMM2.5 (CVCL_C307), OMM2.3 (CVCL_C306) (a gift of Bruce Ksander) and OMM1 (CVCL_6939) were cultured in a mixture of RPMI and DMEM-F12 (1:1) supplemented with 10% FBS and antibiotics [23]. MM66 (CVCL_4D17) was cultured in IMDM supplemented with 20% FBS and antibiotics [24]. The cell lines were maintained in a humidified incubator at 37°C with 5% CO_2_.

### Mass-spectrometry analysis

#### Sample preparation

The experiments were performed according to the published protocol [25].

Briefly, the cells were collected by scraping in lysis buffer, then the lysates were heated to 95°C and cooled on ice, sonicated in a microtip sonicator, heated to 95°C and cooled on ice again. Subsequently, the proteins were precipitated with acetone overnight at -20°C. Protein precipitates were collected by centrifugation, washed with 80% acetone and air dried overnight.

Protein pellets were dissolved in digestion buffer and digested by 1% ProteaseMAX detergent (Promega, Madison, WI, USA) and Trypsin/Lys-C Mix (Promega, Madison, WI, USA) at a 1:50 ratio in a ThermoMixer C (Eppendorf, Nijmegen, The Netherlands) at 2,000 rpm for 18 hours at 37°C.

The digestion was stopped by adding 300 mM KCl, 5 mM KH_2_PO_4_, 50% ACN, 6% Trifluoroacetic acid (TFA), then phosphopeptides were enriched on TiO_2_ beads.

Phosphopeptides were eluted with 40% ACN, 15% NH_4_OH, loaded onto StageTips packed with 3X layers of Empore SPE Disks SDB-RPS material (Sigma-Aldrich St Louis, MO, USA), washed with 0.2% TFA and eluted with 80% ACN, 5% NH_4_OH. The eluates were lyophilized in a freeze-dryer and resuspended in 10 µl 0.1% Formic acid.

#### Mass spectrometry data acquisition

The experiments were performed on an EASY-nLC 1000 system (Proxeon, Odense, Denmark) connected to a Q-Exactive Orbitrap (Thermo Fisher Scientific, Germany) through a nano-electrospray ion source. The Q-Exactive was coupled to a 35 cm analytical column with an inner-diameter of 75 μm, in-house packed with 1.9 μm C18-AQ beads (Reprospher-DE, Pur, Dr. Manish, Ammerbuch-Entringen, Germany) placed into a Butterfly Heater (Phoenix S&T, PA, USA) set to 50°C. The chromatography gradients were performed in 0.1% formic acid increasing the acetonitrile percentage gradually from 5% to 25% acetonitrile in 215 min, then to 30% in 15 min and up to 60% in the next 15 min followed by column re-equilibration. Flow rate was set at 250 nL/min. The mass spectrometer was operated in a Data-Dependent Acquisition (DDA) mode with a top-10 method and a scan range of 300-1600 m/z. Full-scan MS spectra were acquired at a target value of 3 × 10^6^ and a resolution of 70,000, and the Higher-Collisional Dissociation (HCD) tandem mass spectra (MS/MS) were recorded at a target value of 1 × 10^5^ and with a resolution of 17,500, and isolation window of 2.2 m/z, and a normalized collision energy (NCE) of 25%. The minimum AGC target was 1x10^3^. The maximum MS1 and MS2 injection times were 20 and 120 ms, respectively. The precursor ion masses of scanned ions were dynamically excluded (DE) from MS/MS analysis for 60 s. Ions with charge 1, and >6, were excluded from triggering MS2 analysis.

#### Mass spectrometry data analysis

Raw data were analyzed using MaxQuant (version 1.6.2.10) as described previously [26]. We performed the search against an in silico digested UniProt reference proteome for Homo sapiens including canonical and isoform sequences (18^th^ June 2018). Database searches were performed according to standard settings with the following modifications. Oxidation (M), Acetyl (Protein N-term) and Phospho (STY) were allowed as variable modifications with a maximum number of 3. Label-Free Quantification was enabled, not allowing Fast LFQ. Match between runs was performed with 0.7 min match time window and 20 min alignment time window. All peptides were used for protein quantification. All tables were written.

Phospho(STY)sites.txt file from the MaxQuant output was analysed in the Perseus computational platform (1.6.2.2) as described previously [27]. Phosphopeptide intensity values were log_2_ transformed and potential contaminants and proteins identified by site only or reverse peptide were removed. Phosphosites table was expanded into single, double, and multiple phosphosites and filtered to contain quantifications in at least one sample resulting in 14,315 quantified phosphosites. Samples without reasonable depth of sequencing/quality of data (Mel270 replicate 4, OMM2.3 replicate 4 and OMM2.3 replicates 1 and 4) or failed measurements (Mel270 replicate 3) were removed from the total datasets. This quality control step resulted in 13,600 phosphosites and were further subjected to stringent filtering (having intensities in 70% of the samples) resulting in 4,367 phosphosites in total. Missing values were imputed using normally distributed values with a 1.8 downshift (log_2_) and a randomized 0.3 width (log_2_) considering whole matrix values. Two-sided Student’s T-tests with were performed between groups with a cut-off value of p=0.05. Statistical analysis tables were exported and processed in MS Excel, for further filtering and processing of the data. Phosphosites were marked significant when they pass the p-value cut-off during statistical analysis in both metastatic cell lines. The raw data and the analysis can be found in Supplementary Table 1.

### Generation of *MARK3* knock-out cell lines

A Cas9-expressing lentivirus stock was produced by transfecting pKLV2-EF1a-Cas9Bsd-W (Addgene #68343) into HEK293T cells together with packaging vectors (psPax2 and pMD2.G). The MM66 and OMM2.5 cell lines were transduced with this lentivirus and were selected using Blasticidin S. These Cas9-expressing cell lines were subsequently transduced with lentiviruses either expressing a gRNA targeting *MARK3* or a non-targeting gRNA, obtained from the human CRISPR Library (Sigma-Aldrich, St Louis, MO, USA). Transduced cells were selected with puromycin. To generate *MARK3* KO cells we used two distinct gRNAs (sequences: 5’-CACAGCTACATATTTGTTATTGG-3’ (CR-MARK3#1) and 5’-TTTGACTATTTGGTTGCACATGG-3’ (CR-MARK3#2).

Unfortunately, the CR-MARK3#1 gRNA was not efficient in generating a knock-out, and no monoclonal *MARK3* KO cell lines could be generated with this gRNA.

OMM2.3 and OMM1 cells were transduced with the same lentiviruses expressing MARK3-targeting gRNAs and selected with puromycin. Subsequently, these cell lines were transduced with adenovirus vectors expressing either GFP + Cas9, or only GFP as control, as we have described before [28].

As a control for efficiency of generation KO cell lines we used a lentivirus expressing a gRNA targeting the *TP53* gene, described before [28].

### Cell viability assay

The cells were seeded at their appropriate concentrations into clear 96-well plates. The next day, the medium was supplemented with GGTI-298. The treatment was repeated after 2 days. After 5 days from beginning of the experiment, the viability of the cells was assessed using the CellTiter-Blue cell viability assay (Promega, Madison, WI, USA).

### Western blot

The cells were seeded into 6-well plates. Before harvesting, the cells were rinsed 2 times with ice-cold Phosphate Buffered Saline and scraped and lysed with Giordano buffer (50 mM Tris-HCl pH=7.4, 250 mM NaCl, 0.1% Triton X-100, 5 mM EDTA, supplemented with phosphatase- and protease inhibitors. Equal protein amounts were separated on SDS-PAGE and blotted on PVDF membranes (Millipore, Darmstadt, Germany). The membranes were blocked with 10% non-fat dry milk in TBST buffer (10 mM Tris-HCl pH=8.0, 150 mM NaCl, 0.2% Tween-20) and incubated with the primary antibodies diluted in 5% bovine serum albumin/TBST overnight at 4 °C. The membranes were washed with TBST and incubated with horseradish peroxidase-conjugated secondary antibodies (Jackson Laboratories, Bar Harbor, ME, USA). The chemiluminescent signal was visualized using a Chemidoc machine (Biorad, Hercules, CA, USA).

Primary antibodies were obtained from Santa Cruz Biotechnology, Dallas, TX, USA; (MARK3 (F6) and p53 (DO-1), from Cell Signaling Technology, Beverly, MA, USA; (YAP1 (D8H1X) and TAZ (V386), from Sigma-Aldrich, St Louis, MO, USA; (Vinculin (V9131) and from Bethyl Laboratories, Montgomery, TX, USA (USP7; A300-033A).

### RNA isolation and qPCR

The cells were seeded into 6-well plates. The next day, media were supplemented with GGTI-298. After 3 days of treatment, cells were collected by scraping and placed in lysis buffer and RNA was isolated using the SV Total RNA Isolation System (Promega, Madison, WI, USA) according to the manufacturers’ protocol. The reverse transcription reaction was performed using ImPromII reverse transcriptase (Promega). qPCR was performed using SYBR Green Mix (Roche Diagnostics, Indianapolis, IN, USA) in a C1000 Touch Thermal Cycler (Bio-Rad, Hercules, CA, USA). The relative expression of target genes was determined and corrected in relation to the housekeeping genes CAPNS1 and SRPR. In each experiment, the average relative expression was compared to the untreated. Primer sequences are listed in Supplementary Table 2.

### Statistical analysis

The data were analyzed using GraphPad Prism software v.9.1.0 (GraphPad Software, San Diego, CA, USA). Student’s t-test was used to analyze the difference between two groups. One-way ANOVA was used to analyze the differences between multiple groups. P values of 0.05 or less were considered significant.

### Clinical data analysis

The LUMC cohort includes clinical, histopathological, and genetic information on 64 cases treated with primary enucleation at the Leiden University Medical Centre (LUMC) between 1999 and 2008. Clinical information was collected from the Integral Cancer Center West patient records and updated in 2021.

After enucleation, part of the tumor was snap frozen with 2-methyl butane and used for mRNA and DNA isolation, while the remainder was embedded in paraffin after 48 hours of fixation in 4% neutrally buffered formalin and was sent for histological analysis. RNA was isolated with the RNeasy mini kit (Qiagen, Venlo, The Netherlands) and mRNA expression was determined with the HT-12 v4 chip (Illumina, San Diego, CA, USA). Chromosome 3 status was obtained with Single-nucleotide polymorphism analysis, performed with the Affymetrix 250K_NSP-chip and Affymetrix Cytoscan HD chip (Affymetrix, Santa Clara, CA, USA).

TCGA cohort represents 80 primary UM cases enucleated in 6 different centers. mRNA expression was determined by RNA-seq.

The statistical software SPSS, version 25 (IBM Corp, Armonk, NY, USA) was used for statistical analyses of the LUMC and TCGA cohorts. Survival analysis was performed with Kaplan-Meier and log-rank test, with death due to metastases as endpoint. Cases that died of another or unknown cause were censored. The two subpopulations that were compared in each analysis were determined by splitting the total cohort along the median value of mRNA expression for the analyzed gene.

The study was approved by the Biobank Committee of the Leiden University Medical Center (LUMC; 19.062.CBO/uveamelanoomlab-2019-3; B20.023). The tenets of the Declaration of Helsinki were followed.

## Results

### Phosphoproteome analysis of primary and metastatic UM cell lines

To identify the signaling cascades involved in UM metastasis, we performed phosphoproteomics of a primary tumor-derived cell line (Mel270) and two other lines (OMM2.3 and OMM2.5) derived from UM hepatic metastases in quadruplicates. These three cell lines originate from the same individual, share a Q209P mutation in GNAQ, and harbor no mutations in BAP1, EIF1AX or SF3B1. Chromosome analysis of the Mel270 and OMM2.3 lines showed disomy of chromosome 3, tetrasomy of 6p and extra chromosomes 8 [29].

In total, we obtained intensities for 14,315 phosphosites and following stringent quality control step, we quantified 4,367 phosphosites in total. The phosphorylated amino acids were distributed as follows: 3,969 (90.8%) pS, 384 (8.9%) pT and 14 (0.3%) pY. The distribution of phosphorylation multiplicity showed that 2340 (53.6%), 1879 (43%) and 148 (3.4%) sites were phosphorylated at single, double, or multiple sites, respectively. The hierarchical clustering of these 4,367 phosphosites displayed greater separation between primary cell line and the two metastatic cell lines while the two metastatic cell lines also separated well (Fig. 1A).

Comparative analysis uncovered 177 phosphosites to be significantly altered (Student’s T-test, p-value<0.05) between two metastatic UM cell lines (OMM2.3 and OMM2.5) and the primary UM cell line Mel270 (Table 1 and Supplementary Table 1).

**Table 1.** Phosphosites significantly altered between two metastatic UM cell lines (OMM2.3 and OMM2.5) and the primary UM cell line Mel270.

| Gene names_Amino acid Position_Multiplicity | Gene name | -Log Student's T-test p-value omm2.3_mel270 | Log2 (omm2.3/mel270) | -Log Student's T-test p-value omm2.5_mel270 | Log2 (omm2.5/mel270) |
| --- | --- | --- | --- | --- | --- |
| PNN_S100__1 | PNN | 2,320331 | 8,640889 | 2,326058 | 8,241045 |
| ERBB2IP_S932__2 | ERBB2IP | 2,268686 | 8,470394 | 2,958135 | 7,317758 |
| RASAL2_S877__2 | RASAL2 | 3,20149 | 8,333005 | 2,054684 | 6,952386 |
| SRRM2_S1404__2 | SRRM2 | 2,692575 | 8,224896 | 1,868759 | 6,993231 |
| ESCO2_S75__1 | ESCO2 | 2,562178 | 8,201956 | 1,522422 | 6,608265 |
| RASAL2_S880__2 | RASAL2 | 3,90723 | 8,101601 | 2,888066 | 6,720983 |
| CHD1_S1688__1 | CHD1 | 2,308205 | 8,066828 | 2,482959 | 7,221378 |
| TJP1_S605__1 | TJP1 | 3,180118 | 7,891733 | 1,787064 | 6,33797 |
| RRAGC_S95__1 | RRAGC | 3,014953 | 7,501115 | 1,379549 | 6,178071 |
| SPAG9_S194__2 | SPAG9 | 2,425856 | 7,390155 | 1,898788 | 5,164712 |
| XRCC1_T226__2 | XRCC1 | 2,875986 | 7,348891 | 2,017536 | 6,077906 |
| SPAG9_S183__2 | SPAG9 | 3,316868 | 7,330834 | 2,17075 | 6,32826 |
| AHNAK_S5739__1 | AHNAK | 2,524264 | 7,191865 | 1,805079 | 5,962292 |
| LARP1_S90__1 | LARP1 | 2,601892 | 7,160233 | 1,872783 | 6,213362 |
| TMPO_T208__1 | TMPO | 2,896412 | 7,094262 | 2,272455 | 5,42924 |
| GPHN_S127__2 | GPHN | 2,161983 | 6,929839 | 1,471395 | 5,604192 |
| SPAG9_S185__2 | SPAG9 | 2,140266 | 6,712272 | 3,452904 | 6,417998 |
| HSF1_S363__1 | HSF1 | 2,696823 | 6,705507 | 2,782979 | 5,556595 |
| TNS3_S420__1 | TNS3 | 2,589706 | 6,695664 | 1,526324 | 5,546488 |
| FAM83H_S250__1 | FAM83H | 2,496251 | 6,673758 | 1,92563 | 4,863586 |
| PRR12_T1561__2 | PRR12 | 2,581235 | 6,643726 | 1,593389 | 5,458888 |
| PDE4D_S59__2 | PDE4D | 2,479143 | 6,605954 | 1,3098 | 5,063583 |
| TNIK_S740__1 | TNIK | 3,612384 | 6,356614 | 2,168933 | 4,661845 |
| JPH1_S220__2 | JPH1 | 3,185261 | 6,329502 | 2,421806 | 5,579263 |
| EPB41L2_S613__1 | EPB41L2 | 3,686437 | 6,281228 | 3,121418 | 4,778723 |
| RCOR1_S460__1 | RCOR1 | 1,846028 | 6,255202 | 2,562084 | 5,780265 |
| ATXN2_S679__2 | ATXN2 | 1,810305 | 6,198335 | 1,753906 | 5,561078 |
| CDK13_T871__1 | CDK13 | 2,538078 | 6,131416 | 2,029476 | 4,644557 |
| EPB41L2_S612__1 | EPB41L2 | 2,768466 | 6,127415 | 1,689263 | 4,62491 |
| GTSE1_S575__2 | GTSE1 | 2,643891 | 6,125573 | 2,009009 | 4,778836 |
| XRCC1_S235__2 | XRCC1 | 2,655986 | 6,120244 | 1,842754 | 4,849258 |
| CTNND1_S252__1 | CTNND1 | 2,05667 | 6,113639 | 1,788081 | 4,697748 |
| MKI67_T1991__2 | MKI67 | 2,556213 | 6,1061 | 1,366884 | 4,969499 |
| ACTR8_S166__1 | ACTR8 | 2,963651 | 6,054322 | 2,627399 | 5,035097 |
| ZNF318_S1243__1 | ZNF318 | 2,231534 | 6,054263 | 2,051044 | 4,478581 |
| HIRIP3_S370__1 | HIRIP3 | 3,954943 | 6,001266 | 2,222852 | 5,618008 |
| PRR12_S1568__2 | PRR12 | 2,415128 | 5,971622 | 1,460491 | 4,786783 |
| SPAG9_T191__2 | SPAG9 | 2,536629 | 5,963344 | 1,720939 | 4,602517 |
| PRKD1_S548__1 | PRKD1 | 3,116462 | 5,962165 | 2,27624 | 4,249267 |
| TOE1_S5__1 | TOE1 | 1,753938 | 5,954744 | 1,716579 | 4,808053 |
| EGLN1_S125__1 | EGLN1 | 2,857895 | 5,919385 | 1,796896 | 4,478971 |
| JPH1_T448__2 | JPH1 | 2,798863 | 5,902094 | 1,90509 | 4,473262 |
| GTSE1_S580__2 | GTSE1 | 2,505914 | 5,865825 | 1,807325 | 4,519089 |
| MAP3K3_S166__1 | MAP3K3 | 2,312343 | 5,858965 | 1,834668 | 4,289849 |
| TMEM106B_S33__1 | TMEM106B | 3,000869 | 5,754457 | 1,415835 | 3,974044 |
| TCF3_S101__1 | TCF3 | 2,488973 | 5,733511 | 1,591653 | 4,109531 |
| BUB1_S593__2 | BUB1 | 3,375849 | 5,717204 | 1,829598 | 3,570386 |
| MDC1_S485__1 | MDC1 | 2,666944 | 5,679147 | 2,418456 | 4,41726 |
| DST_S2919__1 | DST | 2,806226 | 5,663026 | 1,645212 | 5,231556 |
| TJP1_S912__1 | TJP1 | 1,927121 | 5,640709 | 2,090857 | 4,063324 |

**Table 1.** (continued)
|  |  |  |  |  |  |
| --- | --- | --- | --- | --- | --- |
| STUB1_S23___1 | STUB1 | 2,189052 | 5,633894 | 1,378019 | 3,962584 |
| TNKS1BP1_S1666___1 | TNKS1BP1 | 2,231932 | 5,562864 | 1,416567 | 3,900805 |
| SORBS2_S344___2 | SORBS2 | 2,422821 | 5,492752 | 1,362648 | 4,325774 |
| SORBS2_S346___2 | SORBS2 | 2,422821 | 5,492752 | 1,362648 | 4,325774 |
| BRSK1_S511___1 | BRSK1 | 3,037206 | 5,472331 | 1,64215 | 3,743617 |
| MTM1_S18___1 | MTM1 | 2,4744 | 5,376583 | 1,499966 | 4,001469 |
| JPH1_S452___2 | JPH1 | 2,422273 | 5,364996 | 1,504618 | 3,936164 |
| OSBPL3_S304___1 | OSBPL3 | 2,66361 | 5,353166 | 1,493494 | 3,876441 |
| WRNIP1_S75___1 | WRNIP1 | 2,351422 | 5,338128 | 2,505737 | 3,776667 |
| ZNF318_S1856___1 | ZNF318 | 2,836324 | 5,300036 | 1,848118 | 3,729122 |
| CTTN_S417___2 | CTTN | 2,637005 | 5,284897 | 1,829708 | 3,859484 |
| HIRA_S610___2 | HIRA | 2,44548 | 5,27123 | 1,775098 | 3,811494 |
| PDE4D_S284___1 | PDE4D | 3,176871 | 5,227415 | 1,305763 | 3,549265 |
| KIF21A_S1199___1 | KIF21A | 2,600429 | 5,211912 | 1,464811 | 3,144364 |
| ACIN1_S352___2 | ACIN1 | 2,214204 | 5,152445 | 1,990831 | 3,309524 |
| CAMSAP2_S1121___1 | CAMSAP2 | 1,889824 | 5,150283 | 1,699402 | 3,913262 |
| TCOF1_S171___2 | TCOF1 | 2,133965 | 5,12712 | 1,559842 | 3,779662 |
| PPFIBP1_S447___3 | PPFIBP1 | 2,7788 | 5,096501 | 1,431068 | 2,931659 |
| MAP2_S1782___1 | MAP2 | 1,956307 | 5,068739 | 1,442695 | 4,316577 |
| PRKD1_S549___1 | PRKD1 | 2,500695 | 5,062782 | 1,490412 | 3,349884 |
| TNKS1BP1_S1024___1 | TNKS1BP1 | 3,031735 | 5,020459 | 1,38034 | 3,339321 |
| KIF20A_S514___1 | KIF20A | 2,346576 | 5,014309 | 1,374915 | 3,525265 |
| KIAA0195_S467___1 | KIAA0195 | 2,011465 | 4,972063 | 1,305692 | 4,59376 |
| MAPKBP1_S1075___1 | MAPKBP1 | 2,238285 | 4,905191 | 1,485241 | 3,328155 |
| BAIAP2L2_S451___1 | BAIAP2L2 | 2,981735 | 4,729639 | 1,724977 | 2,91421 |
| HIRA_S612___2 | HIRA | 2,575035 | 4,71989 | 3,71532 | 3,260154 |
| TFIP11_S98___1 | TFIP11 | 1,966688 | 4,70675 | 1,759056 | 3,459704 |
| JPH1_S216___2 | JPH1 | 2,966486 | 4,68522 | 2,547877 | 3,93498 |
| FAM208A_S979___2 | FAM208A | 1,917645 | 4,678004 | 1,429208 | 3,537229 |
| HELZ_S1615___1 | HELZ | 2,910573 | 4,664665 | 2,200548 | 3,688425 |
| LIMA1_S362___1 | LIMA1 | 2,481092 | 4,657305 | 1,918401 | 3,343395 |
| ZNF576_S23___1 | ZNF576 | 2,312478 | 4,641251 | 1,581668 | 3,39615 |
| NAV3_S1190___2 | NAV3 | 1,526135 | 4,618491 | 1,728742 | 3,589816 |
| GTF2I_Y79___1 | GTF2I | 2,861768 | 4,607582 | 1,905497 | 3,825706 |
| INTS10_S28___1 | INTS10 | 1,909621 | 4,539656 | 2,033319 | 3,565192 |
| NAV3_S1189___2 | NAV3 | 1,50071 | 4,486849 | 1,755137 | 3,458174 |
| VCL_S346___1 | VCL | 2,489016 | 4,439525 | 1,456603 | 4,23619 |
| LZTS1_S50___1 | LZTS1 | 1,645176 | 4,439265 | 1,797248 | 2,975595 |
| SORBS2_S417___1 | SORBS2 | 3,192156 | 4,407377 | 1,866019 | 3,067004 |
| ARHGEF11_T1461___2 | ARHGEF11 | 1,333038 | 4,393947 | 1,854821 | 3,925725 |
| TRMT10A_S318___1 | TRMT10A | 2,806635 | 4,352537 | 2,80137 | 3,023454 |
| NME1-NME2_S265___1 | NME1-NME2 | 1,843706 | 4,339633 | 1,855448 | 2,425934 |
| SCRIB_T475___1 | SCRIB | 1,883833 | 4,302811 | 1,389351 | 3,092653 |
| TCP1_S320___1 | TCP1 | 1,869216 | 4,27508 | 1,846498 | 3,023278 |
| MYO5A_S600___1 | MYO5A | 2,281053 | 4,244164 | 1,803709 | 2,702554 |
| MAPT_S232___1 | MAPT | 2,861958 | 4,219584 | 1,52609 | 3,139043 |
| KIAA1109_S4612___1 | KIAA1109 | 2,406284 | 4,175565 | 1,415593 | 2,338774 |
| SPEN_S1278___1 | SPEN | 1,500086 | 4,132838 | 1,732827 | 2,879772 |
| MAP7D1_S86___1 | MAP7D1 | 1,478594 | 4,109335 | 2,190579 | 3,461679 |
| ARHGAP31_S346___1 | ARHGAP31 | 2,44578 | 4,100852 | 1,669664 | 3,386358 |
| ARHGEF7_S516___1 | ARHGEF7 | 2,083972 | 4,086313 | 2,045758 | 2,61045 |
| INCENP_S312___3 | INCENP | 2,38039 | 4,053528 | 1,43104 | 3,280114 |

Table 1. (continued)
|  |  |  |  |  |  |
| --- | --- | --- | --- | --- | --- |
| SNTB1_S389__1 | SNTB1 | 2,198672 | 4,034518 | 1,325791 | 3,046955 |
| DIP2B_S100__1 | DIP2B | 2,109784 | 4,023792 | 1,885077 | 3,181891 |
| NSUN2_S357__1 | NSUN2 | 2,351218 | 4,018664 | 1,329648 | 2,510872 |
| NAV1_S819__1 | NAV1 | 2,942902 | 3,995465 | 1,908395 | 2,495599 |
| RICTOR_S1388__2 | RICTOR | 1,581062 | 3,986675 | 1,755504 | 2,533193 |
| PTPN1_S50__1 | PTPN1 | 1,955495 | 3,978933 | 4,128371 | 1,813383 |
| BAZ1B_S347__1 | BAZ1B | 2,427764 | 3,967728 | 1,522676 | 2,284177 |
| PPFIBP2_S67__1 | PPFIBP2 | 3,112321 | 3,961967 | 1,509483 | 2,271859 |
| CGN_S149__1 | CGN | 2,675491 | 3,873994 | 1,801435 | 2,347765 |
| TCOF1_S906__1 | TCOF1 | 2,888918 | 3,870422 | 1,760603 | 2,50899 |
| MLIP_T135__2 | MLIP | 2,28447 | 3,836281 | 1,682248 | 2,40477 |
| MON1A_S64__1 | MON1A | 2,242929 | 3,767194 | 2,112221 | 1,978688 |
| AHNAK_S5752__1 | AHNAK | 2,225449 | 3,693034 | 2,11725 | 2,551395 |
| ARHGEF2_S196__1 | ARHGEF2 | 3,18585 | 3,648309 | 2,775792 | -3,05644 |
| SIPA1L2_S1488__1 | SIPA1L2 | 2,586717 | 3,590309 | 1,923113 | 1,891082 |
| FOXM1_T711__2 | FOXM1 | 1,988807 | 3,548108 | 1,355429 | 1,59677 |
| CDC25C_S55__2 | CDC25C | 3,076168 | 3,491254 | 1,934797 | 3,041707 |
| LARP7_T344__2 | LARP7 | 1,336955 | 3,478319 | 2,431372 | 1,761458 |
| TJP1_S1487__1 | TJP1 | 2,801101 | 3,466013 | 2,579193 | 2,454255 |
| PHIP_S1783__1 | PHIP | 3,276471 | 3,46228 | 1,459985 | 1,706525 |
| EMD_S173__1 | EMD | 1,70478 | 3,43758 | 1,322032 | -4,77058 |
| ZBTB21_S411__1 | ZBTB21 | 3,425467 | 3,42332 | 2,588625 | 2,249149 |
| HSF1_S303__1 | HSF1 | 1,361852 | 3,41929 | 1,711162 | 2,243472 |
| ANLN_S517__2 | ANLN | 2,246515 | 3,417024 | 2,048682 | 2,798349 |
| ANXA2_S12__1 | ANXA2 | 1,992307 | 3,409278 | 1,351751 | 2,456995 |
| ZNF608_S964__1 | ZNF608 | 1,847191 | 3,388404 | 1,854577 | 3,188241 |
| KIF16B_S662__1 | KIF16B | 1,382869 | 3,321535 | 1,810152 | 1,745818 |
| HMGXB4_S497__1 | HMGXB4 | 2,104139 | 3,319221 | 1,948051 | 1,847479 |
| CASC3_S117__1 | CASC3 | 2,270714 | 3,309143 | 1,646314 | 1,941312 |
| MICALL1_S484__2 | MICALL1 | 1,884212 | 3,302014 | 1,389907 | 1,881289 |
| MICALL1_S486__2 | MICALL1 | 1,884212 | 3,302014 | 1,389907 | 1,881289 |
| CDC25B_S375__1 | CDC25B | 1,354262 | 3,299366 | 3,547596 | 2,638251 |
| ILK_S217__1 | ILK | 1,43567 | 3,250893 | 2,025503 | 1,501854 |
| TNIK_S571__1 | TNIK | 3,051548 | 3,249062 | 1,656601 | 1,935293 |
| HDAC1_S409__1 | HDAC1 | 1,626491 | 3,246864 | 1,562885 | 1,975478 |
| FMN1_S530__1 | FMN1 | 1,51648 | 3,19901 | 1,736315 | 1,576665 |
| TRIOBP_S171__1 | TRIOBP | 1,541126 | 3,168416 | 1,873288 | 0,765635 |
| INCENP_S314__3 | INCENP | 2,324095 | 3,160938 | 1,361574 | 2,387524 |
| NAV1_S103__1 | NAV1 | 1,728778 | 3,061455 | 1,801629 | 1,523159 |
| AHNAK_S5763__1 | AHNAK | 1,985461 | 3,023725 | 1,697325 | -3,41666 |
| DNMT3A_S75__1 | DNMT3A | 1,881033 | 3,001957 | 1,305662 | 2,096782 |
| LIG1_T182__1 | LIG1 | 2,6195 | 2,987185 | 1,575949 | 2,085989 |
| MAPT_S552__2 | MAPT | 1,948161 | 2,863329 | 2,701347 | 1,753795 |
| MAPT_T548__2 | MAPT | 1,948161 | 2,863329 | 2,701347 | 1,753795 |
| COBLL1_T315__2 | COBLL1 | 1,56267 | 2,858164 | 1,591365 | 1,213923 |
| AP3B1_S227__1 | AP3B1 | 1,48636 | 2,698777 | 1,639431 | 1,325895 |
| CCNL1_S335__2 | CCNL1 | 1,692334 | 2,683365 | 1,933779 | -7,1822 |
| EIF4G1_S1046__1 | EIF4G1 | 2,69681 | 2,662518 | 3,280758 | 2,165616 |
| IRS1_S348__1 | IRS1 | 1,385602 | 2,63518 | 1,896005 | 2,087225 |
| BACE2_S415__2 | BACE2 | 1,661837 | 2,620349 | 1,514606 | 1,132684 |
| LATS1_S181__1 | LATS1 | 1,772975 | 2,57788 | 1,394203 | 0,904054 |
| PRKD1_S205__2 | PRKD1 | 1,395023 | 2,359995 | 1,703735 | 1,252979 |

**Table 1.** (continued)
|  |  |  |  |  |  |
| --- | --- | --- | --- | --- | --- |
| PRKD1_S208__2 | PRKD1 | 1,395023 | 2,359995 | 1,703735 | 1,252979 |
| RNF169_T410__3 | RNF169 | 1,414928 | 2,284362 | 1,319031 | 1,194876 |
| SSBP3_T250__1 | SSBP3 | 1,66804 | 2,240761 | 2,252237 | 1,39275 |
| SLC1A5_S317__1 | SLC1A5 | 1,582213 | 2,220436 | 2,874208 | 1,029798 |
| OSBPL3_S309__2 | OSBPL3 | 1,360713 | 2,060791 | 1,393556 | 0,767565 |
| MON2_S205__1 | MON2 | 1,42528 | 1,89273 | 1,726918 | -3,57144 |
| SCRIB_S1220__1 | SCRIB | 1,83174 | 1,860787 | 1,743541 | 0,940604 |
| ZNF106_S1370__1 | ZNF106 | 1,673556 | 1,832644 | 2,062451 | -3,97011 |
| SRGAP1_S936__2 | SRGAP1 | 1,840648 | 1,743917 | 2,510153 | -2,34698 |
| TOP2A_S1391__2 | TOP2A | 1,440786 | 1,690183 | 3,755418 | -4,8608 |
| TOP2A_S1392__2 | TOP2A | 1,440786 | 1,690183 | 1,37965 | 0,681154 |
| TOP2A_S1393__2 | TOP2A | 1,440786 | 1,690183 | 1,37965 | 0,681154 |
| PDE4DIP_S903__1 | PDE4DIP | 2,113536 | 1,66637 | 1,667979 | -0,68882 |
| PDE4DIP_T904__1 | PDE4DIP | 2,113536 | 1,66637 | 1,475524 | -0,7219 |
| PHACTR4_S254__2 | PHACTR4 | 2,655822 | 1,474211 | 1,517245 | 0,584294 |
| PHACTR4_S275__2 | PHACTR4 | 2,655822 | 1,474211 | 1,517245 | 0,584294 |
| MYCBP2_S2749__2 | MYCBP2 | 1,817208 | 0,846 | 2,122912 | -0,50602 |
| MYCBP2_S2751__2 | MYCBP2 | 1,817208 | 0,846 | 2,122912 | -0,50602 |
| TACC2_S56__2 | TACC2 | 2,101844 | -2,3101 | 1,35654 | -3,06379 |
| TACC2_S60__2 | TACC2 | 2,101844 | -2,3101 | 1,35654 | -3,06379 |
| CANX_S475__1 | CANX | 1,827609 | -4,90162 | 2,046562 | -6,52963 |

Pathway analysis on the phosphosites upregulated in UM metastases revealed enrichment of Rho/Rac/Rnd GTPase signaling as well as mRNA splicing and cell cycle pathways (Table 2). GTPase activity is essential for the process of microtubule polymerisation during cell motility and division.

**Table 2.** Pathway analysis on the phosphosites upregulated in UM metastases.

| Pathway name | Entities FDR |
| --- | --- |
| Signaling by Rho GTPases | 2,49E-09 |
| Signaling by Rho GTPases, Miro GTPases and RHOBTB3 | 3,16E-09 |
| RHO GTPase cycle | 2,59E-07 |
| Cell Cycle | 2,12E-04 |
| Cell Cycle, Mitotic | 5,15E-04 |
| Insulin-like Growth Factor-2 mRNA Binding Proteins (IGF2BPs/IMPs/VICKZs) bind RNA | 1,16E-03 |
| Apoptotic execution phase | 2,89E-03 |
| Apoptotic cleavage of cellular proteins | 7,21E-03 |
| Cell Cycle Checkpoints | 1,06E-02 |
| SUMO E3 ligases SUMOylate target proteins | 1,87E-02 |
| RHO GTPases Activate Formins | 1,87E-02 |
| RHOJ GTPase cycle | 1,87E-02 |
| RHOA GTPase cycle | 2,19E-02 |
| SUMOylation | 2,19E-02 |
| RHOQ GTPase cycle | 2,19E-02 |
| Mitotic Anaphase | 2,19E-02 |
| Mitotic Metaphase and Anaphase | 2,19E-02 |
| Amplification of signal from the kinetochores | 2,19E-02 |
| Amplification of signal from unattached kinetochores via a MAD2 inhibitory signal | 2,19E-02 |
| RHOC GTPase cycle | 2,19E-02 |
| M Phase | 2,19E-02 |
| RAC3 GTPase cycle | 3,23E-02 |
| RND3 GTPase cycle | 3,23E-02 |
| Unwinding of DNA | 3,23E-02 |
| RND2 GTPase cycle | 3,23E-02 |
| RHOU GTPase cycle | 3,23E-02 |
| MECP2 regulates neuronal receptors and channels | 3,23E-02 |
| mRNA Splicing - Major Pathway | 3,71E-02 |
| Metabolism of RNA | 3,71E-02 |
| RHOB GTPase cycle | 3,83E-02 |
| CDC42 GTPase cycle | 3,83E-02 |
| Depolymerization of the Nuclear Lamina | 3,95E-02 |
| RAC2 GTPase cycle | 4,05E-02 |
| RHOG GTPase cycle | 4,56E-02 |
| Mitotic Spindle Checkpoint | 4,59E-02 |
| RHO GTPase Effectors | 4,81E-02 |
| mRNA Splicing | 4,86E-02 |

Pathway analysis on the downregulated phosphosites resulted in enrichment of the cascades responsible for immune response, such as interleukin signalling and virus antigen presentation (Table 3).

**Table 3.** Pathway analysis on the phosphosites downregulated in UM metastases.

| Pathway name | Entities FDR |
| --- | --- |
| Maturation of spike protein | 1,03E-02 |
| Assembly of Viral Components at the Budding Site | 1,03E-02 |
| Interleukin-27 signaling | 1,03E-02 |
| Virus Assembly and Release | 1,03E-02 |
| Interleukin-35 Signalling | 1,03E-02 |
| Calnexin/calreticulin cycle | 1,31E-02 |
| Antigen Presentation: Folding, assembly and peptide loading of class I MHC | 1,31E-02 |
| Translation of Structural Proteins | 1,31E-02 |
| N-glycan trimming in the ER and Calnexin/Calreticulin cycle | 1,31E-02 |
| Maturation of spike protein | 1,31E-02 |
| Interleukin-12 family signaling | 1,93E-02 |
| Translation of Structural Proteins | 2,13E-02 |
| Late SARS-CoV-2 Infection Events | 2,64E-02 |
| MHC class II antigen presentation | 2,64E-02 |
| SARS-CoV-1 Infection | 2,64E-02 |
| Influenza Infection | 2,94E-02 |

To link our phosphoproteome datasets to clinical endpoints in UM, we performed correlation analysis of the UM risk factor *BAP1* mRNA expression versus mRNA expression of Microtubule Affinity Regulating Kinase 1-4 (*MARK1-4*), reported upstream regulators of microtubule assembly [30]. We identified *MARK3* as having a weak but significant inverse correlation with mRNA expression of *BAP1* (Fig 1B). Reduction of *BAP1* expression occurs due to inactivating mutations and monosomy on chromosome 3 during UM development, and it known to strongly increase metastatic potential. Hence, reverse correlation with *BAP1* mRNA expression means increased *MARK3* expression could be specific for metastases. Indeed, we found significant difference between *MARK3* expression in UM cases with different chromosome 3 status both in LUMC (Fig. 1C) and TCGA (Fig. 1E) patient cohorts. Moreover, in the LUMC cohort *MARK3* expression correlates with worse prognosis (Fig. 1D); in the TCGA cohort we observe a similar trend, but the difference is not significant.

**Figure 1.**
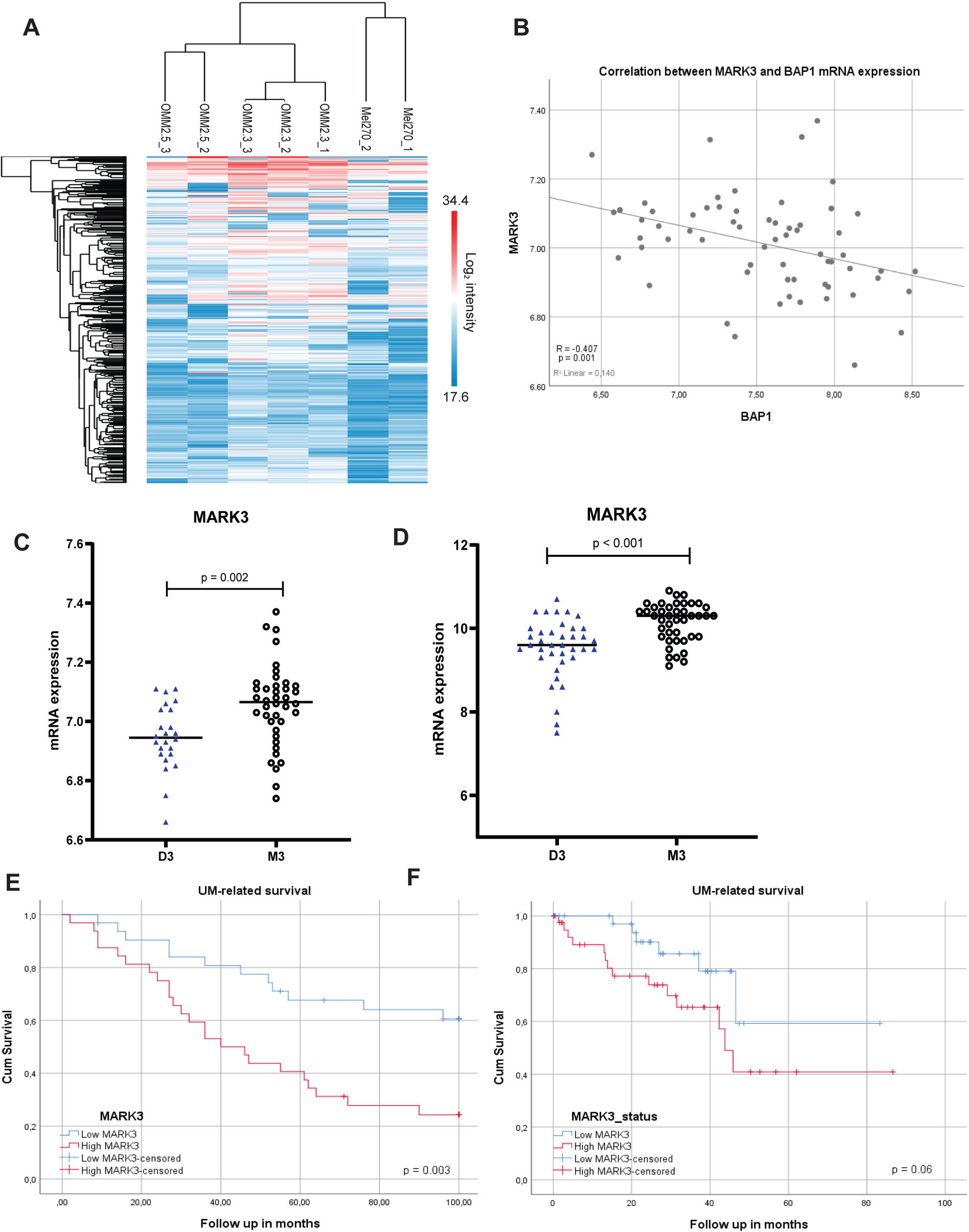
*MARK3* expression correlates with the status of chromosome 3 and survival in UM. (A) Heatmap of the phosphosites identified by mass-spectrometry analysis. (B) Correlation between *MARK3* and *BAP1* mRNA expression in LUMC cohort. (C-D) Correlation of *MARK3* mRNA expression with chromosome 3 status of the tumors in (C) LUMC cohort, (D) TCGA cohort. (E-F) Analysis of the UM-specific survival related to *MARK3* expression in (E) LUMC patient cohort (n=63; split at the median, n=32 high, n=31 low), (F) TCGA cohort, (n=71; split at the median, n=36 high, n=35 low).

We did not find significant correlation between mRNA expression of *MARK1, 2, 4* and any of the tested clinical parameters (data not shown).

MARK3 is known to regulate Rho signaling (top pathway from phosphoproteomics) via ARGHEF2 [31]. Activated Rho/Rac1 are involved in many processes, including an indirect activation/nuclear translocation of transcriptional co-activators YAP1 and TAZ, suggesting a link between MARK3 and nuclear localization of YAP1/TAZ transcription factors via Rho signaling. Since nuclear localization of YAP1/TAZ have been found involved in proliferation of uveal melanoma cells, MARK3 might possibly be important for proliferation or invasion of UM cells.

Taken together, these results indicate potential significance of MARK3-Rho-YAP1/TAZ signaling axis for UM progression. Therefore, we decided to closely study the role of MARK3 on YAP1/TAZ signaling in context of UM.

### Activity of MARK3 might interfere with YAP1/TAZ signaling

In order to evaluate the effect of MARK3 on UM cell viability, we generated MARK3-KO derivatives of UM cell lines MM66, OMM1, OMM2.3 and OMM2.5. In the case of MM66 and OMM2.5, we first introduced a Cas9 lentiviral expression vector and subsequently transduced MM66/Cas9 and OMM2.5/Cas9 with a lentivirus containing either control sgRNA (CR-NT) or sgRNAs targeting *MARK3* (CR-M3) (Suppl. Fig. 1A). In case of OMM1 and OMM2.3 we first stably expressed the *MARK3*-targeting sgRNAs and then transiently introduced Cas9 expressing adenoviral vector or a control vector (expressing GFP) (Suppl. Fig 1B). After generation of these polyclonal cell lines, we isolated monoclonal cell lines lacking MARK3 expression and used these monoclonal cell lines for further investigation. We treated these cell lines with the geranylgeranyl transferase inhibitor GGTI-298, a compound that indirectly attenuates YAP1/TAZ activity by inhibiting the activity of RhoA/Rac1 [32]. As illustrated in Fig. 2A and 2B, depletion of MARK3 in MM66 and OMM2.3 cells slightly reduced the level of TAZ compared to control cells and this effect is enhanced by GGTI-298 treatment. The level of YAP1, however, stays stable across all the conditions, although in OMM2.3 cells the band of YAP1 is migrating slightly slower upon GGTI-298 treatment, which might possibly indicate increased phosphorylation of YAP1. The same effect is illustrated on additional clones of OMM2.3 CR-M3 in Suppl. Fig. 2A.

In OMM1 cells (Fig. 2B) the levels of both YAP1 and TAZ do not change upon MARK3 knockout and GGTI-298 treatment; in OMM2.5 CR-M3 clones TAZ levels are slightly downregulated upon GGTI-298 treatment comparing to vehicle treated samples (Suppl. Fig. 2B).

**Figure 2.**
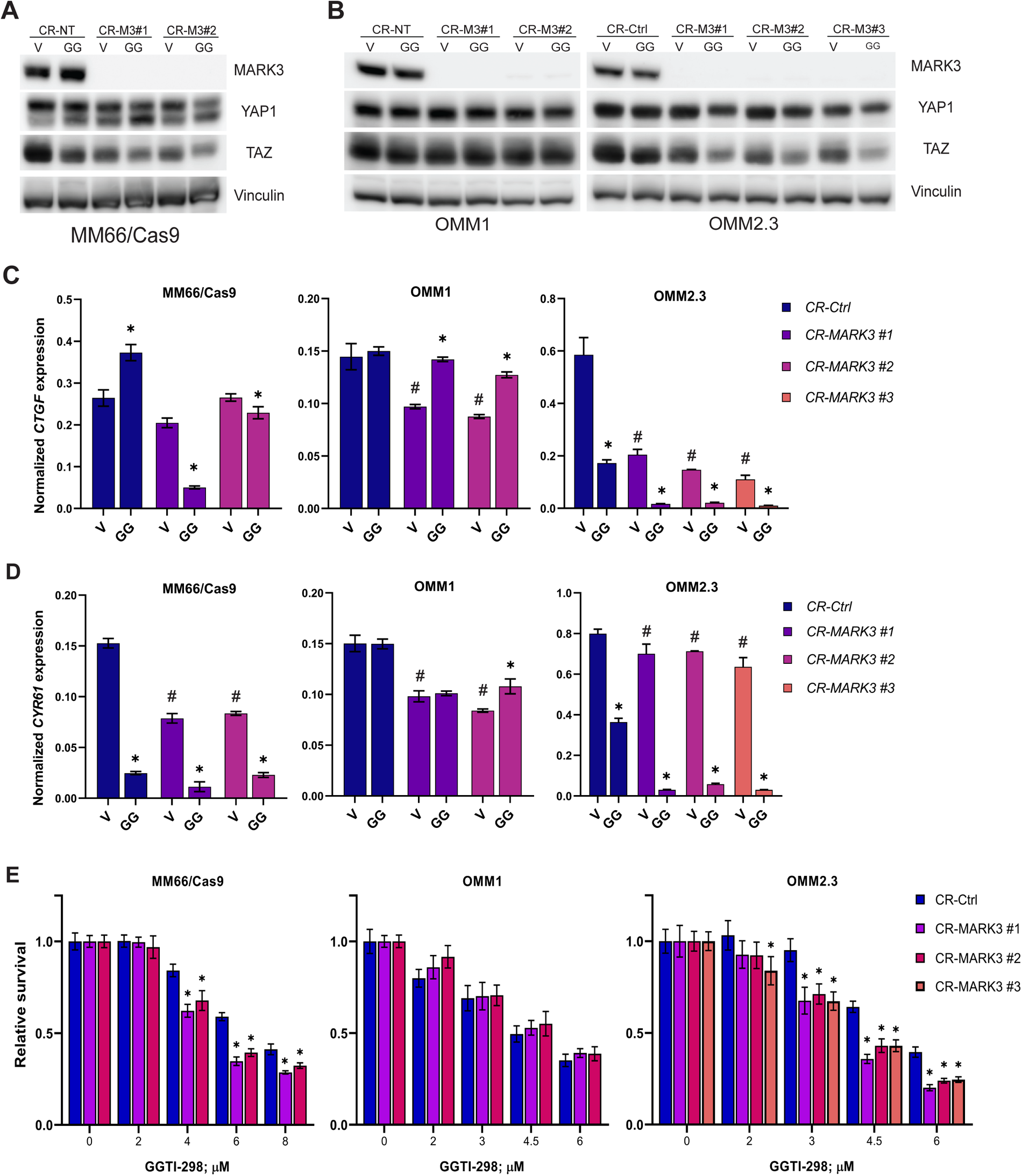
MARK3 knockout in combination with GGTI-298 synergistically inhibits growth of UM cell lines. (A-B) Effect of MARK3 knockout on YAP1 and TAZ protein expression in (A) MM66, (B) OMM1 and OMM2.3. V-vehicle, GG-GGTI-298 (6 µM for MM66 and OMM2.3, 4 µM for OMM1), CR-NT - Non-targeting control, CR-M3 - MARK3 knockout; Vinculin was used as a loading control. (C-D) Expression of (C) *CTGF* and (D) *CYR61* mRNA upon 24h treatment with GGTI-298 in MARK3-knockout UM cell lines. V-vehicle, GG-GGTI-298 (6 µM for MM66 and OMM2.3, 4 µM for OMM1); significant (p<0.05) change in mRNA expression upon MARK3 knockout (*CR-MARK3*) compared to the vehicle *CR-Ctrl* is indicated with (#); significant (p<0.05) change in mRNA expression upon GGTI-298 treatment compared to the vehicle control is indicated with (*), statistical analysis was performed using one-way ANOVA, error bars present mean ± SEM, n=3. (E) Effect of GGTI-298 on viability of MARK3-knockout UM cell lines after 5 days of treatment. Significant (p<0.05) reduction of viability in CR-MARK3 comparing to CR-Ctrl is indicated with (*), statistical analysis was performed using one-way ANOVA, error bars present mean ± SEM, n=3.

To determine whether *MARK3* knockout has an effect on YAP1/TAZ activity, we investigated the expression of two classic YAP1/TAZ target genes *CTGF* (Fig. 2C and Suppl. Fig. 2C) and *CYR61* (Fig. 2D and Suppl. Fig. 2D), both basal and upon GGTI-298 treatment. The basal expression of *CTGF* was reduced in most *MARK3* knockout cell lines compared to the controls, although not consistent in OMM2.5. The basal expression of *CYR61* is not significantly lower in most *MARK3*-KO cell lines, with exception in MM66 cells. In MM66 and OMM2.3 cell lines, treatment with GGTI-298 further downregulates expression of both YAP1/TAZ target genes, but OMM1 demonstrates no, or even the opposite effect. Again, the effects in OMM2.5 cells are not consistent.

Subsequently we investigated the consequence of *MARK3* knock-out on cell viability upon GGTI-298 treatment. Interestingly, the combination of *MARK3* knockout and GGTI-298 treatment synergistically inhibited growth of MM66, OMM2.3 and OMM2.5 cell lines, but not in the OMM1 cell lines (Fig. 2E and Suppl. Fig. E, F). This effect to some extent correlates with the effects of *MARK3* knock-out on expression of the YAP1/TAZ target genes. It is important to note that *MARK3* knockout did not consistently affect the growth rate of the UM cell lines (data not shown).

## Discussion

To detect signaling pathways, involved in UM metastatic spread, we compared phosphoproteomes of the cell lines derived from UM primary tumor and metastases. Mass spectrometry analysis indicated 177 differently phosphorylated sites, and some of the hits: ARHGEF2 [31], TNIK1 [33] HSF1 [34], SORBS2 [35], have been reported to participate in YAP1/TAZ signaling cascade. These results and our previous work [36] indicate the involvement of YAP1/TAZ signaling in the process of UM metastatic spread.

In line with the studies of various cancer types [37], our pathway analysis indicated enrichment of GTPase activity related processes in metastatic cell lines compared to primary tumor. Specifically in UM, the elevated RhoC GTPase activity was reported in the tumors with higher metastatic potential harboring monosomy on chromosome 3 [38].

Interestingly, the down-regulated pathways in metastatic UM cell lines compared to a primary cell line are mostly related to immune response and antigen presentation. This effect has been described in metastatic UM and might be important for immune evasion [39].

MARK3 kinase has been shown to phosphorylate ARHGEF2 and thus stimulate activation of RhoA, which is an essential regulator of YAP1 activity in UM [12]. We demonstrate that full depletion of MARK3 expression results in down regulation of YAP1/TAZ target genes and, in case of MM66 and OMM2.3 cell lines affects the protein levels of TAZ. The inconsistent effect of *MARK3* KO on mRNA expression of CTGF and CYR61 in OMM2.5 might be attributed to possible off-target effects of sgRNAs, since the effects of GGTI-298 treatment is similar to the other cell lines. Moreover, we have previously shown [36] that expression of CYR61 in OMM2.5 is dependent on TAZ, but not YAP1.

We have demonstrated that combination of YAP1/TAZ depletion with the geranylgeranyl transferase inhibitor GGTI-298, acting downstream in the mevalonate pathway and reducing the activity of Rho proteins, synergistically slows down growth of UM cell lines [36]. Similarly, when *MARK3* knockout is combined with GGTI-298, the effect on transcription of YAP1/TAZ target genes is significantly enhanced, and the growth of some *MARK3* KO UM cell lines is synergistically inhibited. However, the synergistic effect of the combination is not very strong and a proportion of the *MARK3* KO cells remain viable even after prolonged incubation with relatively high concentrations of GGTI-298, what can indicate potential activation of resistant mechanisms.

The role of MARK3 in YAP1/TAZ signaling and tumor progression might be context dependent, as follows from the report of Machino et al., which showed that lower *MARK3* expression significantly correlated with poor prognosis in HGSOC patients [40]. This report is in contrast with our finding that high levels of MARK3 correlates with worse prognosis of UM patients and that higher *MARK3* expression correlated with monosomy of chromosome 3.

On the other hand, treatment of glioma cell lines with a recently described MARK3/MARK4 inhibitor reduced their proliferation *in vitro* and tumorigenic growth in a xenograft mouse model [41].

In UM we have not observed the consistent effect of MARK3 knockout on cell viability, and the combination of MARK3 knockout with GGTI-298 was not able to completely abrogate the growth of metastatic UM cell lines.

We conclude that MARK3 appears to be involved in YAP1/TAZ signaling regulation in UM, but its suitability as a therapeutic target needs further investigation.

## Acknowledgements

We thank dr. Bharath Sampadi for providing the protocol for phosphopeptide enrichment, analysis of the mass spectrometry data and editing the manuscript.

## Conflict of interests

None declared.

## Funding

This research was funded by European Union’s Horizon 2020 project “UM Cure 2020” (grant no. 667787).

## Notes

### Competing Interest Statement

The authors have declared no competing interest.

### Summary of Updates

One extra reference has been added to the Discussion section

