## Supplementary Table 2 for "Comparative phosphoproteome analysis of primary and metastatic uveal melanoma cell lines"

| Primer | Sequence |
|---|---|
| CAPNS1 FW | ATGGTTTTGGCATTGACACATG |
| CAPNS1 RV | GCTTGCCTGTGGTGTCGC |
| SRPR FW | CATTGCTTTTGCACGTAACCAA |
| SRPR RV | ATTGTCTTGCATGCGGCC |
| CTGF FW | GTTTGGCCCAGACCCAACTA |
| CTGF RV | GGCTCTGCTTCTCTAGCCTG |
| CYR61 FW | CAGGACTGTGAAGATGCGGT |
| CYR61 RV | GCCTGTAGAAGGGAAACGCT |

**Supplementary Table 2.** Primers for qPCR.
