## Supplementary material for "Comparative phosphoproteome analysis of primary and metastatic uveal melanoma cell lines": Suppl. Fig. 1

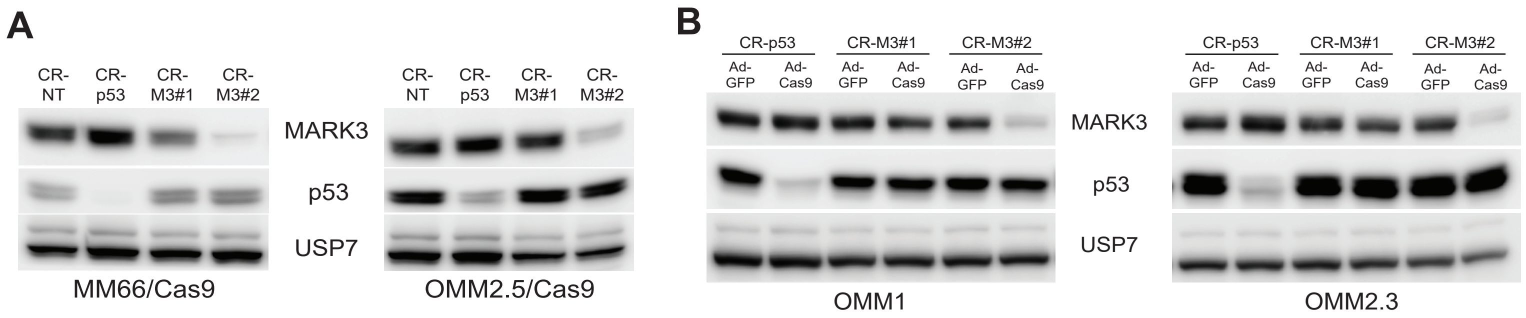


**Supplementary Figure 1. Kinase enrichment analysis and generation of MARK3 knockout derivatives of UM cell lines**
(A) MM66 and OMM2.5 cell lines were transduced with a lentiviral vector expressing Cas9 and subsequently transduced with a lentivirus containing either control sgRNA (CR-NT) or sgRNAs targeting *MARK3* (CR-M3).
(B) OMM1 and OMM2.3 were transduced with a lentiviral vector expressing *MARK3*-targeting sgRNAs (CR-M3) and then transiently introduced Cas9 expressing adenoviral vector or a control vector (expressing GFP).
sgRNA targeting *p53* (CR-p53) was used as a positive control for Cas9 activity, USP7 -loading control.
