## Supplementary material for "Comparative phosphoproteome analysis of primary and metastatic uveal melanoma cell lines": Suppl. Fig. 2

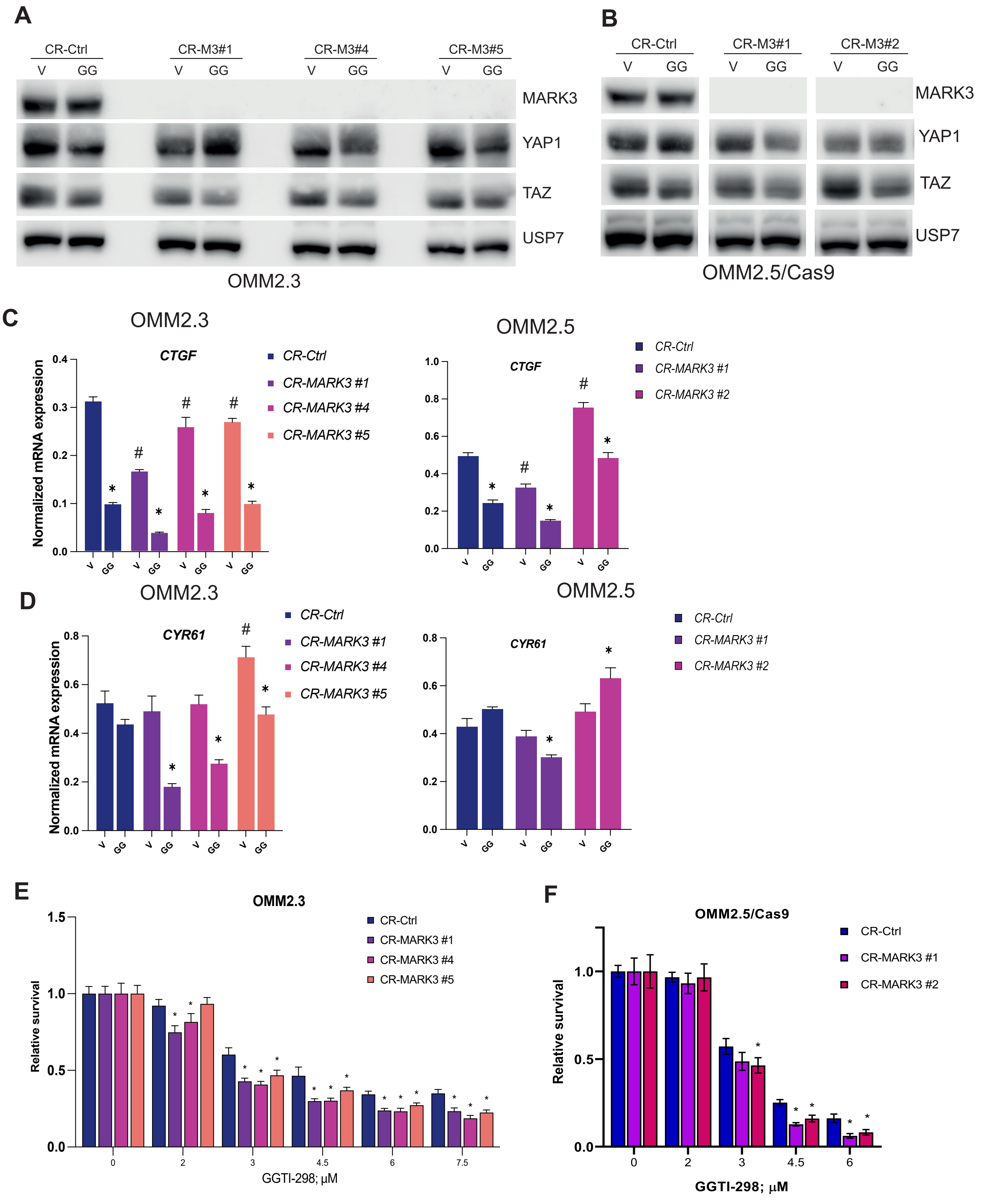


**Supplementary Figure 2. Effect of MARK3 knockout on YAP1/TAZ signaling**

(A-B) Effect of MARK3 knockout on YAP1 and TAZ protein expression in (A) OMM2.3, (B) OMM2.5. V-vehicle, GG-GGTI-298 (6 µM), CR-Ctrl - Non-targeting control, CR-M3 - MARK3 knockout ; USP7 was used as a loading control.

(C-D) Expression of (C) *CTGF* and (D) *CYR61* mRNA upon 24h treatment with GGTI-298 in MARK3-depleted UM cell lines. V-vehicle, GG-GGTI-298 (6 µM); significant ($p<0.05$) change in mRNA expression upon MARK3 knockout (CR-MARK3) compared to the vehicle control (CR-Ctrl) is indicated with (#); significant ($p<0.05$) change in mRNA expression upon GGTI-298 treatment compared to the vehicle control is indicated with (*), statistical analysis was performed using one-way ANOVA, error bars present mean ± SEM, n=3.

(E) Effect of GGTI-298 on viability of MARK3-depleted UM cell lines after 5 days of treatment. Significant ($p<0.05$) reduction of viability in MARK3 knockout (CR-MARK3) comparing to control (CR-Ctrl) is indicated with (*), statistical analysis was performed using one-way ANOVA, error bars present mean ± SEM, n=3.
